# Whole genome sequence-based characterization of foodborne *Staphylococcus aureus* isolated from pork in 2018 and 2023

**DOI:** 10.1101/2025.06.20.660668

**Authors:** Jiali Zhu, Taya Tang, Linli Ji, Heng Li

**Affiliations:** Department of Microbiology, School of Basic Medical Sciences, Suzhou Medical College, Soochow University, Suzhou 215123, China

**Keywords:** Foodborne pathogens, Whole-genome sequencing, Antimicrobial resistance, Virulence factors, One Health

## Abstract

This study characterizes the genomic epidemiology of 111 *Staphylococcus aureus* isolates collected from retail meat in Beijing and Copenhagen during 2018 and 2023 using whole-genome sequencing. Our analysis identified concerning antimicrobial resistance patterns, with methicillin-resistant *S. aureus* (MRSA) prevalence rising from 19.51% to 24.13% over the study period. The livestock-associated CC398 lineage (27.03% of isolates) demonstrated strong correlations with tetracycline resistance (*tetM*) and persisted as the dominant clone in Beijing, increasing from 33.33% to 40.91% prevalence. In contrast, we observed the rapid emergence of community-associated CC8 strains in Denmark, reaching 71.43% prevalence by 2023^27^. Virulence profiling revealed MSSA strains frequently carried enterotoxin genes (*seg*/*sei* in 55.8% of isolates), while mobile genetic elements like SCCmec IV in ST59-t437 contributed significantly to pathogenicity. Phylogenetic analysis delineated five major clades and highlighted the expansion of multidrug-resistant CC398 strains in Beijing. These findings demonstrate the dynamic evolution of *S. aureus* at the human-animal interface and emphasize the urgent need for integrated One Health surveillance systems to track resistance gene dissemination. Future research should prioritize investigating zoonotic transmission pathways and the role of horizontal gene transfer in driving the convergence of virulence and resistance traits in foodborne pathogens.

## Introduction

*Staphylococcus aureus* (*S. aureus*) poses a substantial public health threat due to its widespread presence in both food products and human populations^15^.^19^ This pathogen frequently contaminates raw meat, dairy products, and aquatic foods, primarily through inadequate hygiene practices during processing or cross-handling, often leading to acute foodborne illnesses^1^. From 2010 to 2020, China documented 17,985 cases of foodborne illnesses, predominantly attributed to fungal and bacterial pathogens including *S. aureus*^3^. Similarly, *S. aureus* was responsible for 10.4% of reported foodborne disease cases from 2007 and 2011 in Europe^2^, highlighting its global significance as a causative agent of foodborne outbreaks. These findings emphasized the urgent need for improved food safety protocols and heightened regulatory oversight to reduce associated public health risks.

As the conditional bacteria of foodborne risks, methicillin-resistant *Staphylococcus aureus* (MRSA) persisted as a critical global public health concern due to its multidrug resistance and high pathogenicity^20^, despite its declining incidence worldwide^4^.^10^ However, methicillin-sensitive *Staphylococcus aureus* (MSSA), which constituted the majority of clinical infections (e.g., 79% of pediatric cases in the U.S.), remained epidemiologically significant^5^. MSSA serves as a key reservoir for antimicrobial resistance dissemination, capable of rapid conversion to MRSA through *SCCmec* acquisition^16^.

As a foodborne pathogen, *S. aureus* produced an array of virulence factors, including α-hemolysin (*hla*), Panton-Valentine leukocidin (PVL), staphylococcal enterotoxins (SEs; e.g., *sea*, *seb*), and toxic shock syndrome toxin-1 (*tsst*-1), in addition to biofilm formation mediated by the *icaAD* operon^13^. Among these, SEs represented a primary causative agent of staphylococcal food poisoning, highlighting the significant public health threat posed by this bacterium^14^.

In 2018 and 2023, retail pork samples were collected in Beijing, China and Copenhagen, Denmark, for isolation of *S. aureus* strains, followed by whole-genome sequencing (WGS). This investigation characterized the genetic diversity, plasmid profiles, antibiotic resistance patterns, and virulence determinants of *S. aureus* across these geographically distinct regions. The resulting genomic data offered critical insights to inform evidence-based interventions for mitigating foodborne illnesses associated with this pathogen.

## Materials and methods

### Bacterial isolation and identification

A total of 248 retail pork samples were purchased in Beijing and Copenhagen from 2018 to 2023. Briefly, 10 g of meat were homogenized in 0.1% peptone saline, serially diluted, and plated onto Baird Parker agar (HopeBio 4115, Beijing, China) and CHROMagar™ MRSA agar (Becton Dickinson, Franklin Lakes, NJ). The plates were incubated overnight at 37°C in a CO₂ incubator and examined for S. aureus colonies. A total of 111 *Staphylococcus aureus* strains were subsequently confirmed using MALDI-TOF MS (BioMérieux, France).

### Whole-genome sequencing (WGS)

The isolates were cultured in Tryptone Soya Broth (AOBOX, China) at 37°C for 24 hours. Genomic DNA was extracted using the HiPure Bacterial DNA Kit (Meiji Biotech, China), and purity and concentration were measured using a Nanodrop ND-1000 spectrophotometer. Whole-genome sequencing was conducted on the Illumina NextSeq 500 platform^11^ (Honsunbio, China). Raw sequencing reads were assembled and assessed with EToKi v1.0, and QUAST v2.3.

### Molecular typing and phylogenetic analysis

Sequence types (STs) were identified using MLST 2.0, and clonal complexes (CCs) were assigned using eBURST v3^22^. For phylogenetic analysis, a maximum-likelihood tree was constructed with CSI Phylogeny v1.4, using S. aureus ATCC 25923 as the reference strain.

### Identification of mobile genetic elements, antimicrobial resistance and virulence genes

Antimicrobial resistance (AMR) genes, virulence factors, and mobile genetic elements were detected using ResFinder, VFDB, and MobileElementFinder, respectively^21^. For each gene, the minimum thresholds were set at ≥60% sequence coverage and ≥90% sequence identity for all databases, except MobileElementFinder which required ≥60% identity for detection.

### Statistical analysis and visualization

The phylogenetic relationships were plotted with Grapetree and iTOL. The line chart and donut chart were made using GraphPad Prism 7, and statistical significance was assessed using One-way ANOVA with p < 0.05.

## Results

### Prevalence of staphylococcus aureus

This study analyzed 111 *Staphylococcus aureus* isolates collected from Beijing and Denmark with 82 isolates (15 from Beijing, 67 from Denmark) obtained in 2015 and 29 isolates (22 from Beijing, 7 from Denmark) collected in 2023. Among these, we identified 23 methicillin-resistant *S. aureus* (MRSA) strains, yielding an overall prevalence of 20.72% (23/111)^29^. The MRSA prevalence of the two cities increased from 19.51% (16/82) in 2015 to 24.13% (7/29) in 2023.

### Sequence typing and clonal distribution

Among the 19 clonal complex (CC) types identified, CC398 (n=30, 25.64%) emerged as the predominant lineage, followed by CC8 (n=9), CC7 (n=9), CC5 (n=8), and others. MRSA strains were predominantly clustered within the CC398 (16/30), CC8 (3/9), and CC59 (2/2) lineages (**Table 1**). *Spa* typing revealed substantial diversity, with over 40 distinct spa types detected. The most frequent subtypes were t034 (n=18) and t008 (n=8). CC1 contained t127, t273, and t3324 (t273/t3324 predominant), while CC398 exhibited the greatest diversity (t011, t034, t571) with t034 showing a transmission advantage (60%, n=18). Among 111 isolates, MLST identified 24 sequence types (STs) classified by eBURST v3 into 17 CCs, dominated by CC398 (27.03%, n=30), particularly in Danish 2015 isolates (n=16), followed by widely distributed CC5 (ST5, 7.21%, n=8) and CC7 (ST7, 8.11%, n=9), along with CC8 (8.11%)^23^.

**Table 1.**
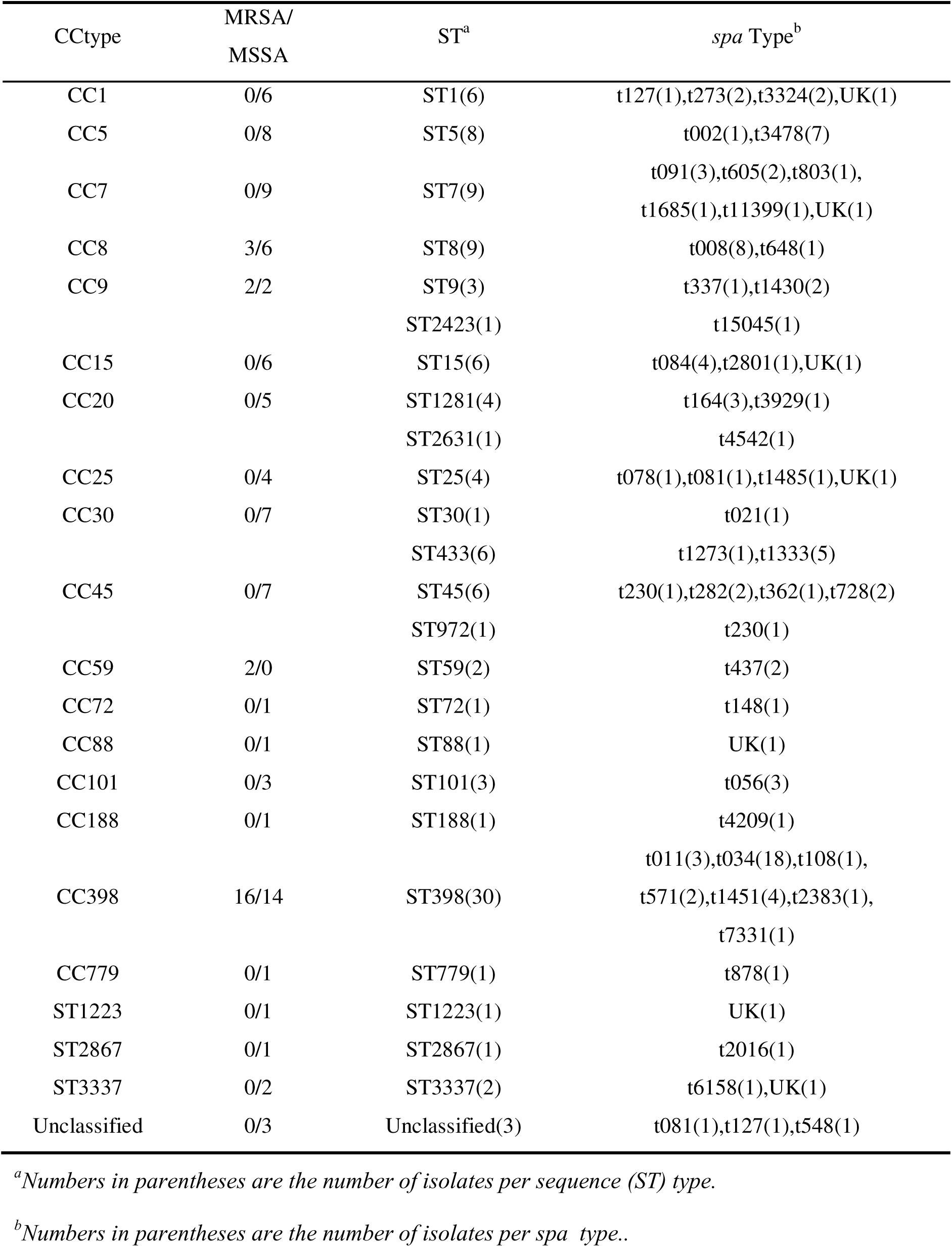
CC, MRSA/MSSA, ST, and *spa* type of *S. aureus* isolates from this study.

### Genetic characteristics of MRSA and MSSA

The 23 MRSA strains were exclusively restricted to ST8, ST9, ST59, and ST398 lineages, while MSSA isolates (n=88) demonstrated greater genetic diversity across multiple sequence types (**Table 1**). MRSA distribution followed distinct clonal complex (CC)-specific patterns. The CC398 predominated (69.57%, 16/23), likely reflecting its adaptation to hospital or livestock-associated environments. CC8 and CC9 contained 3 and 2 MRSA strains, respectively, with their ST8 (t008) and ST9 (t1430) variants potentially representing community-acquired MRSA (CA-MRSA). Notably, CC59 consisted exclusively of MRSA strains (2/2), and its ST59 (t437) lineage may be associated with stable resistance determinants (e.g., SCCmec type IV). In contrast, MSSA isolates (n=88) were distributed across 16 CCs, primarily CC398 (n=14), CC5 (n=8), and CC7 (n=9).

### Plasmid replicons and insertion sequences

Among the 111 *S. aureus* strains analyzed, 13 plasmid replicon types (rep-types) and 3 insertion sequences (ISSau) were identified. The predominant replicon was rep16 (n=46, 39.3%), which was widely distributed across lineages such as CC398, CC1, and CC7. Other common replicons included rep5a (n=33), rep7a (n=23), and rep7c (n=23). Notably, 63.3% of CC398 strains (19/30) carried rep16, along with enrichment of ISSau1 and ISSau3. In contrast, MSSA-associated lineages (e.g., CC15 and CC30) exhibited a lower prevalence of these mobile genetic elements (**Table 2**).

**Table 2.** Distribution of various rep-types in *Staphylococcus aureus* strains in this study.

| CC types | No.<sup>a</sup> | rep5a<br>(n=33) | rep7a<br>(n=23) | rep7c<br>(n=23) | rep10<br>(n=14) | rep16<br>(n=46) | rep19<br>(n=23) | rep20<br>(n=13) | rep21<br>(n=25) | repUS43<br>(n=18) | ISSau1<br>(n=18) | ISSau3<br>(n=16) | ISSau5<br>(n=17) |
| --- | --- | --- | --- | --- | --- | --- | --- | --- | --- | --- | --- | --- | --- |
| CC1 | 6 | 2 | 0 | 6 | 1 | 4 | 2 | 0 | 3 | 1 | 0 | 0 | 0 |
| CC5 | 8 | 0 | 4 | 0 | 1 | 0 | 0 | 1 | 4 | 1 | 0 | 0 | 7 |
| CC7 | 9 | 8 | 5 | 0 | 1 | 3 | 0 | 5 | 0 | 0 | 0 | 3 | 0 |
| CC8 | 9 | 0 | 1 | 9 | 0 | 3 | 3 | 5 | 3 | 0 | 0 | 0 | 3 |
| CC9 | 4 | 0 | 0 | 0 | 0 | 2 | 0 | 0 | 2 | 0 | 0 | 3 | 0 |
| CC15 | 6 | 5 | 0 | 0 | 0 | 4 | 1 | 2 | 3 | 0 | 0 | 0 | 0 |
| CC20 | 5 | 0 | 0 | 0 | 4 | 0 | 0 | 0 | 0 | 0 | 0 | 0 | 0 |
| CC25 | 4 | 0 | 0 | 4 | 0 | 2 | 2 | 0 | 0 | 0 | 0 | 0 | 0 |
| CC30 | 7 | 1 | 1 | 0 | 0 | 1 | 0 | 0 | 1 | 0 | 7 | 0 | 1 |
| CC45 | 7 | 1 | 0 | 0 | 0 | 2 | 0 | 0 | 1 | 0 | 0 | 7 | 0 |
| CC59 | 2 | 0 | 2 | 0 | 0 | 2 | 2 | 0 | 0 | 0 | 0 | 0 | 0 |
| CC72 | 1 | 1 | 0 | 1 | 1 | 1 | 0 | 0 | 0 | 0 | 0 | 1 | 0 |
| CC88 | 1 | 0 | 0 | 0 | 0 | 0 | 0 | 0 | 0 | 0 | 0 | 0 | 0 |
| CC101 | 3 | 0 | 0 | 0 | 0 | 0 | 0 | 0 | 0 | 0 | 0 | 0 | 3 |
| CC188 | 1 | 1 | 0 | 0 | 0 | 1 | 1 | 0 | 0 | 0 | 0 | 1 | 1 |
| CC398 | 30 | 12 | 10 | 0 | 5 | 19 | 12 | 0 | 8 | 16 | 11 | 0 | 0 |
| CC779 | 1 | 0 | 0 | 0 | 0 | 0 | 0 | 0 | 0 | 0 | 0 | 0 | 0 |
| ST1223 | 1 | 1 | 0 | 0 | 0 | 1 | 0 | 0 | 0 | 0 | 0 | 0 | 0 |
| ST2867 | 1 | 0 | 0 | 1 | 0 | 0 | 0 | 0 | 0 | 0 | 0 | 0 | 0 |
| ST3337 | 2 | 0 | 0 | 0 | 0 | 0 | 0 | 0 | 0 | 0 | 0 | 1 | 2 |
| Unclassified | 3 | 1 | 0 | 2 | 1 | 1 | 0 | 0 | 0 | 0 | 0 | 0 | 0 |

<sup>a</sup>Numbers in parentheses are the number of isolates.

### Distribution of antimicrobial resistance genes

Antimicrobial resistance profiling of 111 *S. aureus* strains identified genes conferring resistance to seven antimicrobial classes, including β-lactams (*blaZ*, *mecA*), aminoglycosides (*aac*, *aph*, *aadD*), tetracyclines (*tetK*, *tetM*), and macrolides (*ermB*, *ermC*). All MRSA strains (23/23) carried *mecA*, with 69.57% (16/23) exhibiting multidrug resistance (≥3 resistance classes), while only 18.18% (16/88) of MSSA strains showed MDR^24^. Resistance patterns varied by lineage that 93.3% (28/30) of CC398 strains harbored *blaZ* and 53.3% (16/30) carried *mecA*, while both CC59 strains (2/2) displayed complete (100%) multidrug resistance (**Table 3**).

**Table 3.**
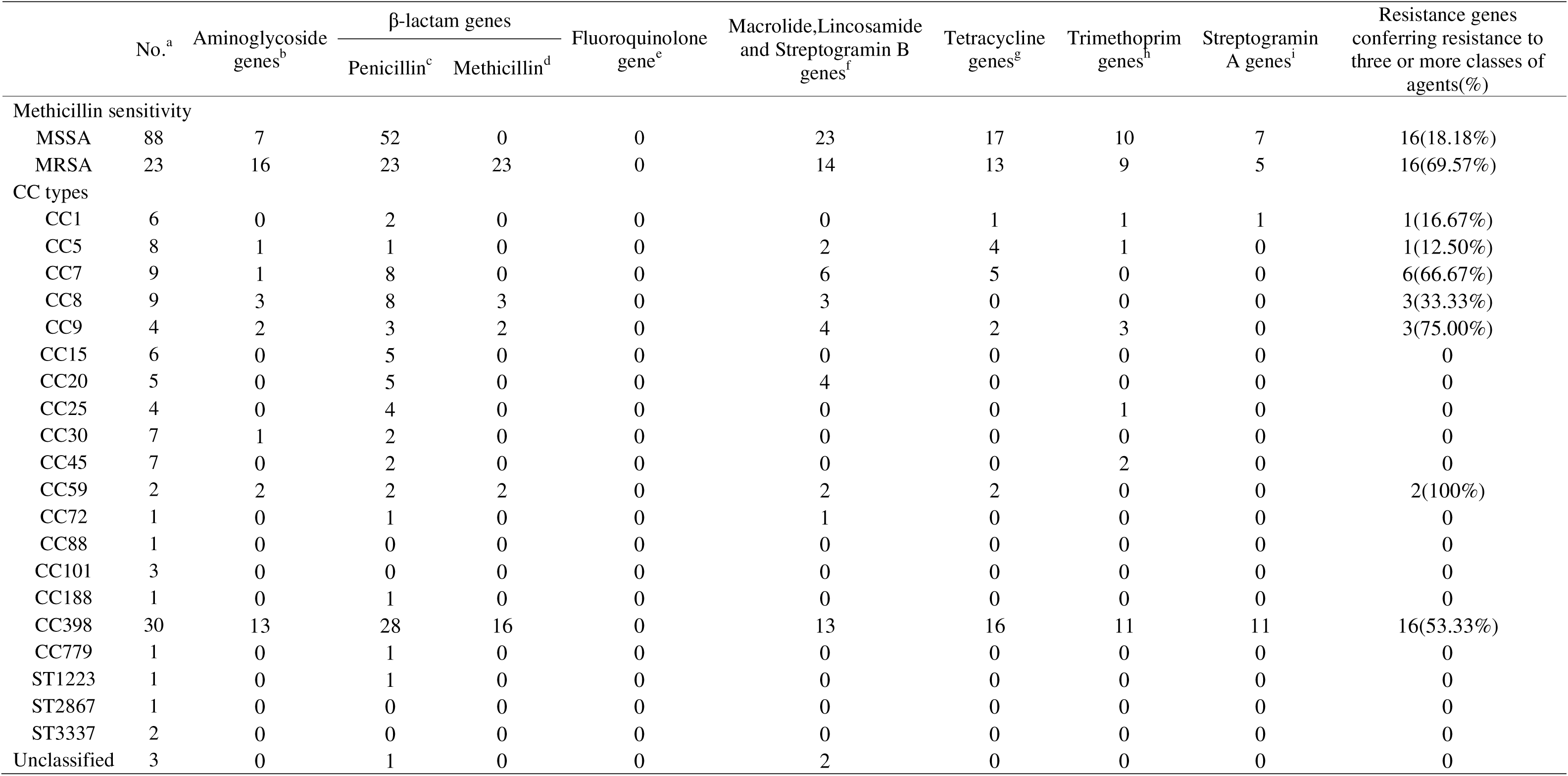
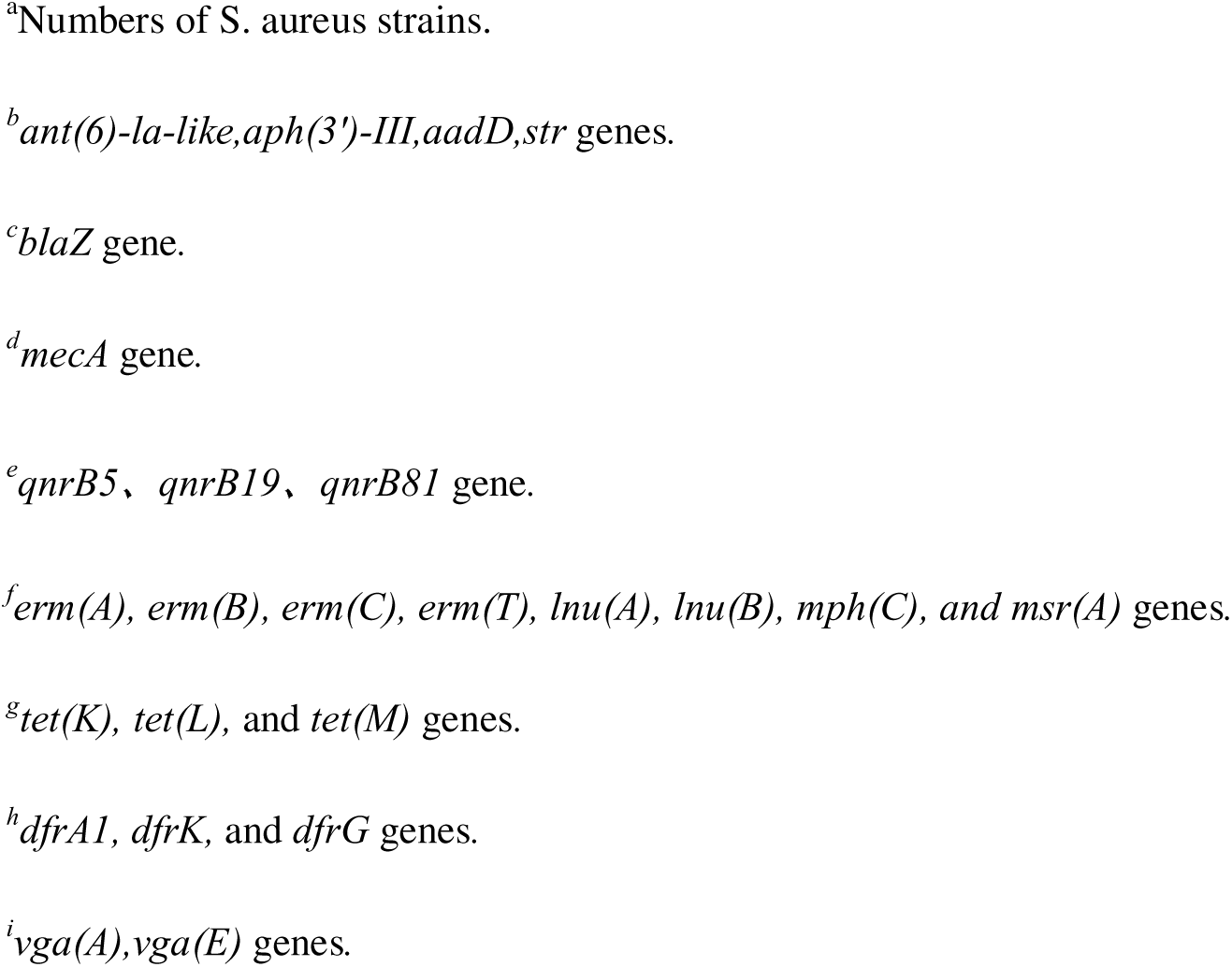
Distribution of antimicrobial resistance genes in *Staphylococcus aureus* strains in this study.

### Patterns of virulence factors

Analysis of 15 virulence factors revealed distinct distribution patterns among the 111 *S. aureus* strains. Hemolysin genes (*hlgA*/*B*/*C*) demonstrated remarkable conservation, present in 95-97% (111-114) of isolates. In contrast, PVL toxin showed limited prevalence (2.56%, 3 strains), exclusively in CC8 and CC30 MRSA isolates. The *tst* toxin occurred in 5.98% (7 strains), primarily within CC398 and CC7 lineages. Serine protease genes (*splA/B/E*) exhibited moderate prevalence (51-57 strains) with strong MSSA lineage association. These findings demonstrate significant lineage-specific distribution of virulence factors, with certain toxins showing preferential absence in MRSA backgrounds (**Table 4**).

**Table 4.**
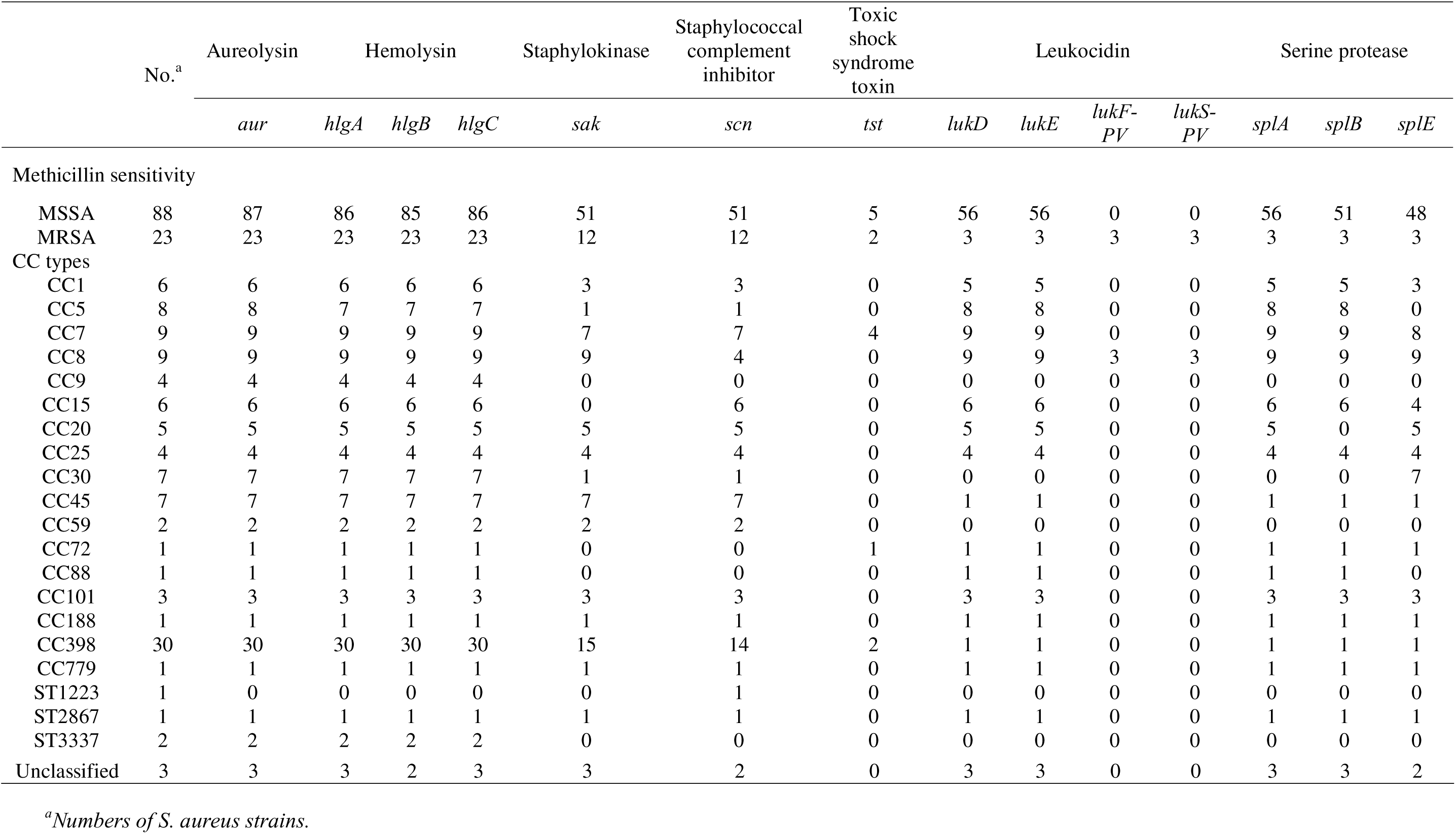
Distribution of virulence factors in *Staphylococcus aureus* strains in this study.

### Lineage-specific enterotoxin gene profiles

Among the 18 enterotoxin genes detected, MSSA strains exhibited significantly higher prevalence (69.32%, 61/88) compared to MRSA (30.43%, 7/23). Type II enterotoxins (*seg*, *sei*, *sem*, *sen*) were ubiquitous in MSSA lineages (CC5, CC30, CC25; 100% positivity), but rare in CC398 (6.67%, 2/30), suggesting this lineage primarily employs non-enterotoxin virulence strategies. Classical enterotoxins (*seb*, *sek*, *sel*) were predominantly restricted to CC8 and CC59. These findings demonstrate distinct lineage-specific retention patterns of enterotoxin genes in these foodborne *S. aureus* strains (**Table 5**).

**Table 5.**
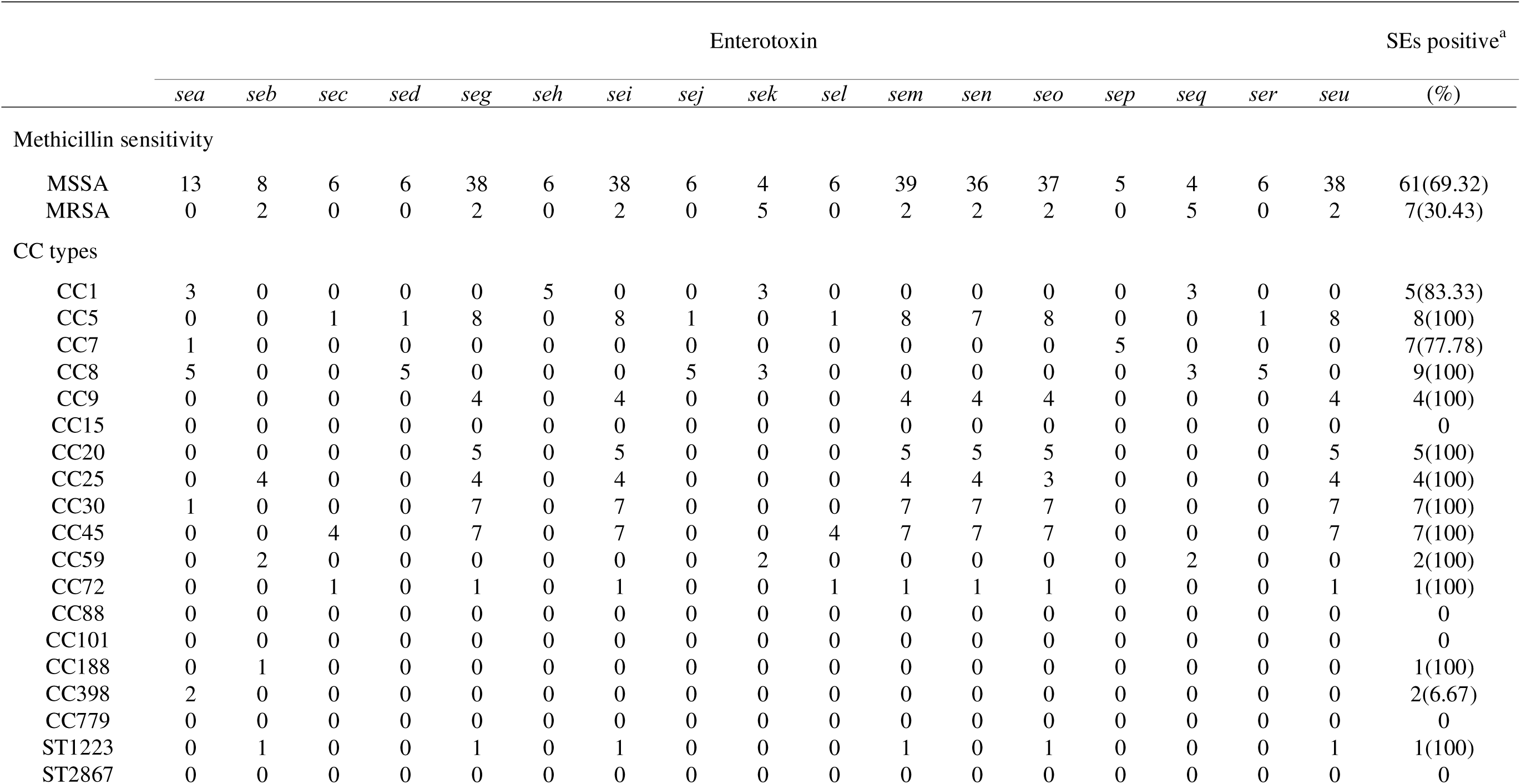
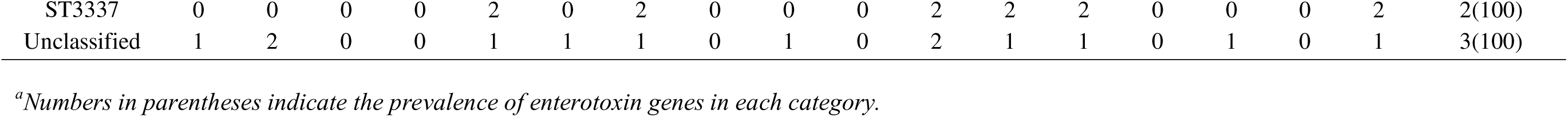
Distribution of enterotoxin genes in *Staphylococcus aureus* strains in this study.

### Clonal complex shifts between Beijing and Copenhagen

Core genome phylogenetics revealed distinct geographical and temporal patterns in CC prevalence (**Figure 1**). In Beijing, CC398 dominated both sampling periods (2015: 33.33%; 2023: 40.91%), with secondary lineages shifting from CC25 (20.00%) and CC59 (13.33%) in 2015 to CC7 (13.64%) and CC20 (9.09%) in 2023. This persistent CC398 dominance correlated with the carriage of livestock-associated spa types (t034, t011), and a 95% *tetM* co-occurrence in farm-derived t034 strains, reflecting selective pressures from regional intensive pig farming.

**Figure 1.**
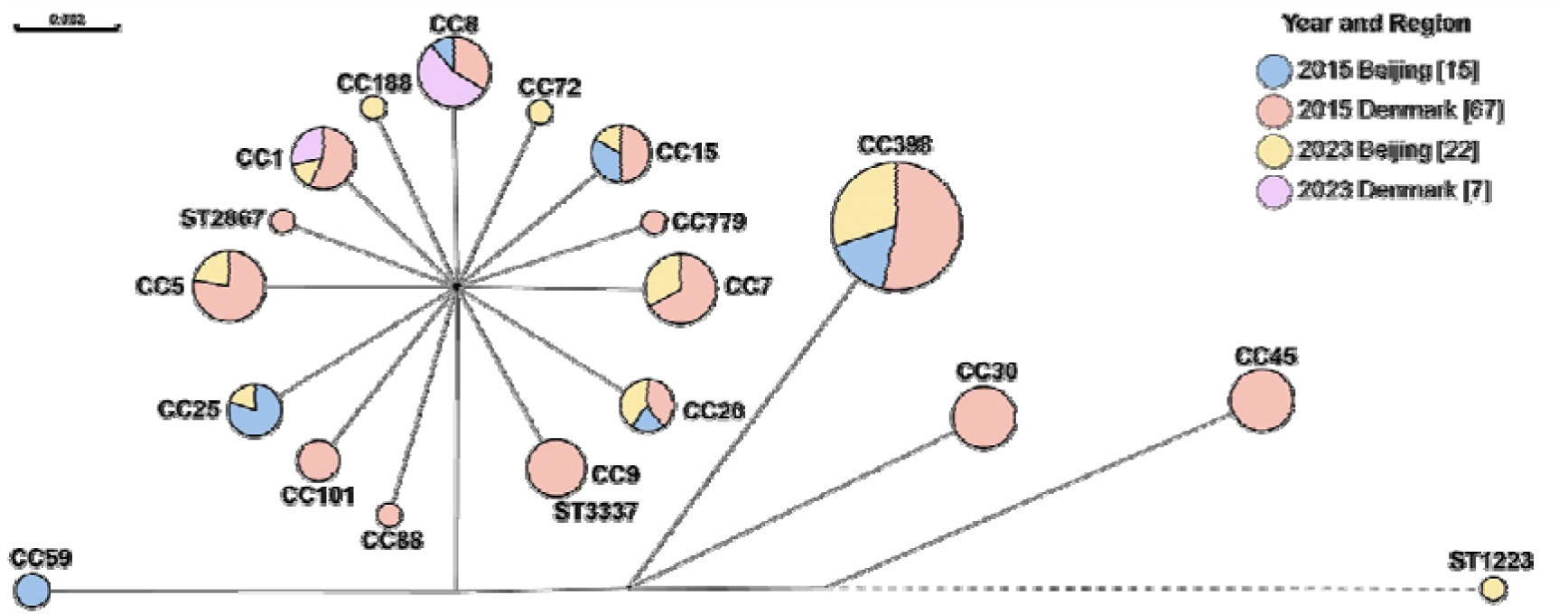
Minimal spanning tree based on the multi-locus sequence types of 111 *S.aureus* isolates collected from East China. Each circle represents a sequence type, circle sizes represent the number of isolates, circle areas are colored by isolation years, and the length of the line connecting the circles represents the relative distance, signifying the degree of genetic variation.

The emerging Beijing lineages (CC7/CC20) carried *icaAD* biofilm genes, suggesting potential hospital-acquired transmission routes. In Denmark, CC398 maintained substantial prevalence (2015: 23.88%) alongside CC30/CC45 (10.45% each), consistent with historical livestock-associated patterns in Nordic regions. The 2023 CC8 surge (27.03%, vs 4.48% in 2015) indicated new community transmission risks, contrasting with Beijing’s stable CC398 epidemiology.

### Phylogenetic analysis and spatiotemporal dynamics

Phylogenetic analysis of 111 *S. aureus* isolates, incorporating spa typing, clonal complex classification, temporal data, and resistance profiles, revealed five distinct evolutionary clades (A-E, **Figure 2/3**). The phylogenetic structure demonstrated phenotypic segregation, with Clade A predominantly comprising hospital-associated MRSA lineages (CC5, CC8, CC45) and Clade B representing livestock-associated MRSA (CC398, CC9). Clade C emerged as a distinct branch containing the Asian epidemic CC59 lineage, notable for its dual carriage of virulence and resistance determinants. In contrast, Clades D and E were primarily composed of diverse MSSA lineages (CC7, CC15, CC25, CC30), forming separate phylogenetic clusters.

**Figure 2.**
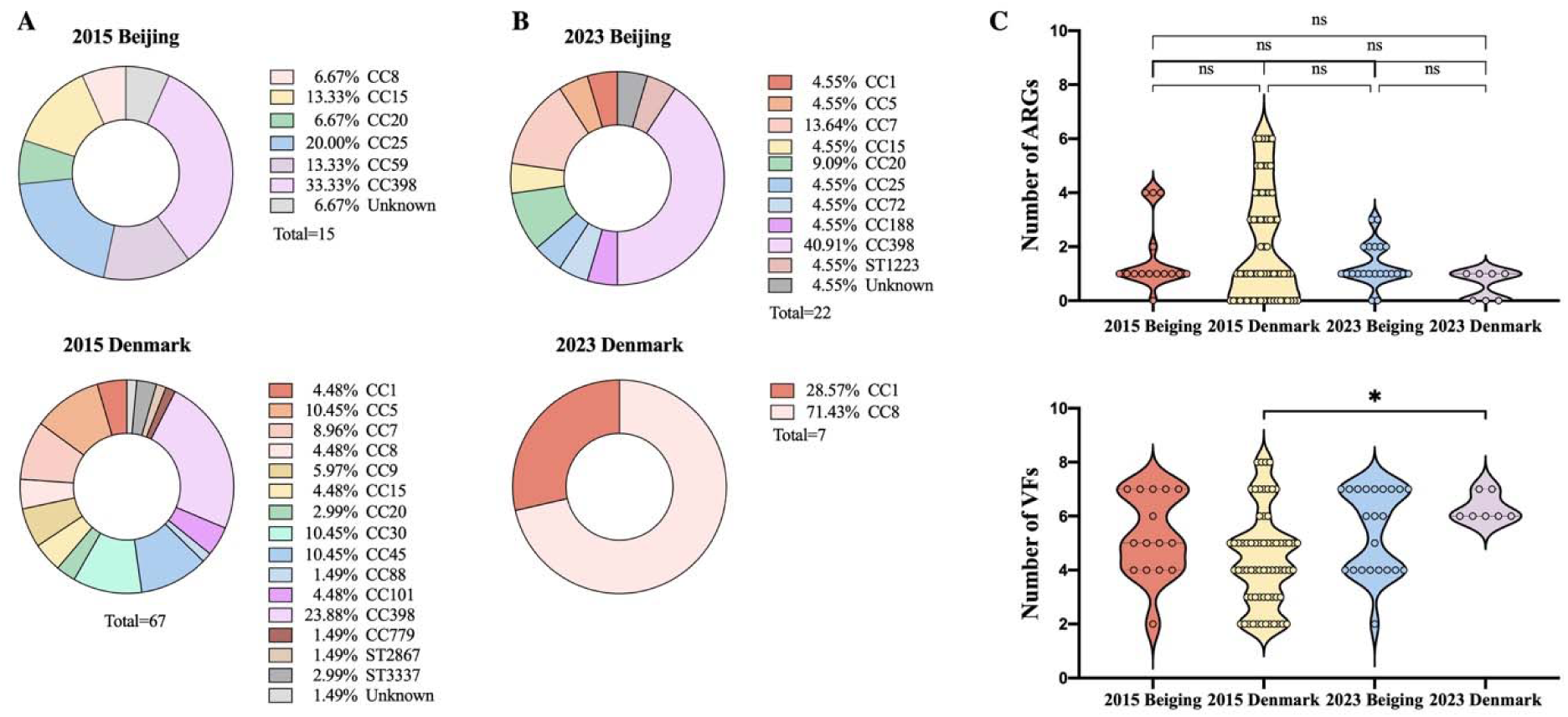
Distribution of CC types and the number of ARGs and VFs of *S.aureus* strains isolated from eastern China between 2019 to 2024. (A-C) Donut charts are colored by clonal comple types. (D, E) Violin Plots represents the number of antimicrobial resistance genes and Virulence factors of *S. aureus* strains in different years, each point represents a single strain.

**Figure 3.**
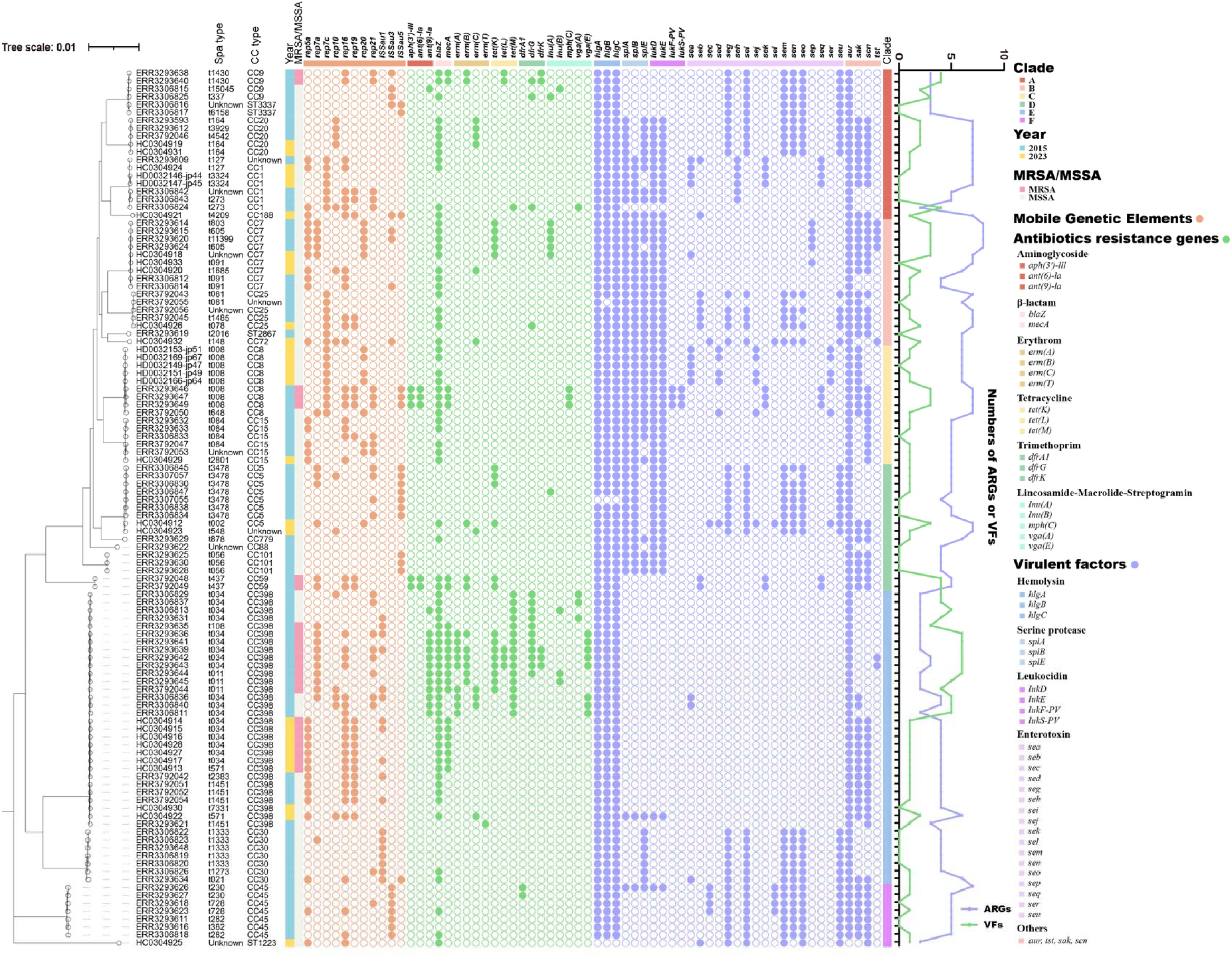
Phylogenetic tree and Molecular characteristics of 106 *S.aureus* isolates. The phylogenetic tree is shown on the left. The squares colored by CC types, isolation year, MRSA/MSSA, MGEs, AMRs, and VFs. The line chart showed the numbers of ARGs and VFs.

Mobile genetic element analysis revealed fundamental differences in horizontal gene transfer potential across clades (**Figure 3**). Clades A and B, encompassing both HA-MRSA and LA-MRSA populations, were enriched for diverse plasmid replicons (rep-types) and insertion sequences (ISSau family), reflecting their enhanced capacity for genetic exchange. This contrasted sharply with the MSSA-dominated Clades D and E, which carried fewer mobile elements and demonstrated more stable genomic architectures.

Distribution patterns of antimicrobial resistance gene closely mirrored phylogenetic divisions (**Figure 2**). The LA-MRSA strains of Clade B consistently carried a characteristic multidrug resistance arsenal including *mecA*, *blaZ*, *tetM* and *tetK*, while HA-MRSA strains in Clade A maintained a distinct resistance profile centered on *mecA*, *aac(6’)-aph(2’’)*, and *erm(B)*. Both MRSA-dominated clades exhibited significantly higher resistance gene burdens compared to the MSSA-predominant Clades D and E, which showed minimal resistance gene carriage.

The virulence factor spectrum displayed both conserved and lineage-specific patterns (**Figure 3**). The CC59 strains in Clade C stood out for their unique combination of multiple resistance mechanisms and virulence factors, representing a true “high virulence-high resistance” phenotype. PVL toxin (*lukF/S-PV*) demonstrated restricted distribution, primarily found in Clade A (HA-MRSA) and Clade E (MSSA) isolates. Enterotoxin genes (*seg, sei, sem, sen*) were particularly abundant in MSSA lineages of Clades D and E (CC5, CC25, CC30), while showing marked depletion in most MRSA lineages.

## Discussion

This study provided critical insights into the genomic epidemiology of *S. aureus* strains isolated from food sources in Beijing and Copenhagen, with implications for public health surveillance, antimicrobial resistance (AMR) control, and food safety policy development.

The data revealed an overall MRSA prevalence of 20.72%, with Denmark exhibiting higher rates (27.03%) compared to Beijing. These findings corroborated previous European reports documenting persistent MRSA circulation in both livestock and community settings^26^. The observed increase from 19.51% in 2015 to 24.13% in 2023 suggested continuing MRSA expansion, potentially attributable to selective pressures from agricultural antibiotic use. Notably, CC398 predominance in Denmark (23.88% in 2015) paralleled reports from other Nordic countries where this lineage associates with swine production systems^17^. In contrast, the low but rising rate of MRSA infection in Beijing may reflect regional differences in antibiotic management or livestock management practices^18^.^30^

The CC398 lineage demonstrated remarkable predominance (27.03%), with the t034 spa type representing 60% of isolates, confirming its status as an established livestock-associated (LA) clone. The strong correlation with tetM (95% co-occurrence) highlighted tetracycline application in swine production as a driver of resistance. Particularly noteworthy is the increasing CC398 prevalence in Beijing, rising from 33.33% to 40.91% between 2015 and 2023, which mirrored the expansion of intensive farming practices in China and underscored the urgency for enhanced veterinary antibiotic regulation^6^.^25^

The detection of CC8 as a potential CA-MRSA clone (71.43% in Denmark, 2023) raised significant concern^7^, particularly given its well-documented association with community-acquired outbreaks^9^. The complete absence of CC8 MRSA isolates in Beijing indicated distinct epidemiological drivers, potentially attributable to differences in healthcare-associated transmission dynamics or host immune profiles.

Comparative genomic analysis revealed fundamental differences between MRSA and MSSA populations. MRSA strains predominantly carried characteristic resistance determinants, exemplified by *SCCmec* IV in CC59 isolates, whereas MSSA variants displayed greater heterogeneity in virulence gene content. Notably, enterotoxin genes (*seg*, *sei*) were identified in 55.8% of MSSA isolates, underscoring the often-overlooked pathogenic potential of methicillin-susceptible variants in foodborne illness. The distribution of enterotoxins genes followed distinct clonal patterns, with CC5 strains harboring *sea* and *seb*, and CC59 isolates carrying *seb*, which both genotypes being epidemiologically linked to staphylococcal food poisoning outbreaks. Surprisingly, CC398 isolates consistently lacked enterotoxin genes, a finding congruent with their established livestock-associated MRSA phenotype and suggestive of evolutionary adaptation favoring colonization over acute virulence expression.

This study also revealed critical differences in mobile genetic element acquisition among *S. aureus* lineages^28^. CC398 MRSA strains exhibited particularly efficient horizontal gene transfer, with 63.3% carrying rep16 plasmids alongside frequent ISSau1 and ISSau3 insertions, suggesting enhanced genomic plasticity. In contrast, MSSA-associated lineages like CC15 and CC30 showed markedly lower mobile element acquisition rates. These findings demonstrate the genetic exchange capacity of MRSA facilitates the co-transfer of resistance and virulence determinants, driving pathogen evolution.

The comprehensive genomic analysis of Beijing and Denmark isolates uncovered distinct geographical patterns in virulence factor distribution, highlighting the complex interplay between bacterial genetics and local ecological pressures. Particularly noteworthy is the persistent dominance of livestock-adapted CC398 strains in Beijing, carrying characteristic resistance markers like tetM, compared to Denmark’s emerging community-associated CC8 clones. These results underscore the necessity of integrated One Health strategies that bridge human medicine, veterinary practice, and food production systems to effectively monitor and contain antimicrobial resistance dissemination^8^.^12^

Several methodological limitations should be acknowledged in this study. The relatively small Danish sample from 2023 (n=7) reduced statistical robustness, while the lack of accompanying clinical data, particularly regarding patient outcomes, precludes definitive conclusions about zoonotic transmission pathways. To address these constraints, subsequent investigations would benefit from expanded sample collection incorporating clinical human isolates, thereby enabling comprehensive One Health analyses across human-animal-environment interfaces.

## Conclusion

This study characterized foodborne *Staphylococcus aureus* strains isolated from foods in Beijing and Copenhagen, highlighting key antimicrobial resistance (AMR) and virulence trends. The overall MRSA prevalence was 20.72% (23/111), with notable geographic variation in Denmark (27.03%) compared to Beijing, a rising trend over time (from 19.51% in 2015 to 24.13% in 2023). The dominant lineage, CC398 (27.03%), was strongly associated with livestock exposure and tetracycline resistance (*tetM*). In contrast, the rapid emergence of CC8 (reaching 71.43% in Denmark by 2023) suggests a growing risk of community-associated MRSA. Key virulence determinants (*seg*/*sei* in 55.8% of MSSA) and mobile genetic elements (SCCmec IV in ST59-t437) were identified as major contributors to pathogenicity. Phylogenetic analysis revealed the expansion of CC398 in Beijing (increasing from 33.33% to 40.91%), likely driven by multidrug resistance. These findings emphasize the urgent need for integrated One Health interventions to control *S. aureus* dissemination across food production, healthcare, and livestock sectors. Future research should investigate zoonotic transmission risks and the role of horizontal gene transfer in virulence and resistance evolution.

## Ethical approval

Not applicable

## Informed Consent Statement

Not applicable

## Consent for publication

Not applicable

## Availability of data and materials

The raw reads from this study are submitted to the NCBI with the accession numbers shown in Supplementary materials.

## Authors’ contributions

JZ, YG, XK, HL, ZZ, XW wrote the main manuscript text and CG, LJ prepared the figures and tables. All authors reviewed the manuscript.

## Acknowledgements

Not applicable

